# A data-driven approach to the automated mapping of functional brain topographies across species

**DOI:** 10.1101/412114

**Authors:** Wolfgang M. Pauli

## Abstract

Behavioral neuroscience has made great strides in developing animal models of human behavior and psychiatric disorders. Animal models allow for the formulation of hypotheses regarding the mechanisms underlying psychiatric disorders, and the opportunity to test these hypotheses using procedures that are too invasive for human participants. However, recent scientific reviews have highlighted the low success rate of translating results from animal models into clinical interventions in humans. A potential roadblock is that bidirectional functional mappings between the human and rodent brain are incomplete. To narrow this gap, we created a framework, Neurobabel, for performing large-scale automated synthesis of human neuroimaging data and behavioral neuroscience data. By leveraging the semantics of how researchers within each field describe their studies, this framework enables region to region mapping of brain regions across species, as well as cross-species mapping of psychological functions. As a proof of concept, we utilize the framework to create a functional cross-species mapping between the amygdala and hippocampus for fear-related and spatial memories, respectively. We then proceed to address two open questions in the field: (1) Do rodents have a dorsolateral prefrontal cortex? (2) Which human brain region corresponds to the rodent prelimbic cortex?

---

Current understanding of the neurobiological processes underlying brain disorders is strongly informed by research in non-human animals, where mechanistic hypotheses can be tested with procedures too invasive for human volunteers. For example, decades of empirical work in rodents have enabled a detailed understanding of the neurobiological mechanisms underlying drug-related behavior in animals (for recent reviews, see [1, 2]). Developing robust, animal-informed models is critical to efforts to treat and diagnose substance abuse disorders, which impose enormous social costs (estimated at between 0.01% and 0.5% of GDP in Europe [3], and about 4% of GDP in the U.S.A., including costs related to crime, lost work productivity and health care [4]).

Though rodent models are a critical component of the evidence base informing theory refinement and treatment innovation, uncertainty about whether analogous neural mechanisms are conserved within the human brain has undermined progress in identifying specific biological mechanisms, and in developing corresponding therapeutic targets in humans [5]. Improving the precision for translating behavioral neuroscience findings into psychiatric interventions is critical, as research aimed at developing new treatments for brain disorders is under threat; prominent pharmaceutical companies have cut research funding dedicated to developing drugs for various psychiatric disorders [6].

The central aim of the present work was therefore to develop a novel data-driven interdisciplinary approach that enables researchers to systematically compare functional topographies across species. To this end, we developed an extension, Neurobabel, for an existing framework for brain mapping, the Neurosynth project [7]. Neurosynth performs large-scale automated synthesis of human neuroimaging data by combining information about the location of reported brain activation with the frequency of scientific terms in each published neuroimaging study (known as term frequency–inverse document frequency, or tf-idf). This process yields a term-to-coordinate mapping that has been highly successful in synthesizing results from neuroimaging studies, by enabling large-scale automated meta-analyses and the decoding of mental states [8, 9, 10, 11].

We build on this existing framework by incorporating data from published rodent studies (N = 2668), and use this augmented dataset to enable a bidirectional functional mapping between the rodent and the human brain. Coordinates of surgical procedures in the rodent brain were extracted from each study and cross-referenced against the frequency of scientific terms in the companion articles. The resulting dataset can be probed to synthesize findings across rodent studies, analogous to the functionality of Neurosynth for human neuroimaging studies. By using the distribution of word frequencies across a vocabulary of scientific terms shared by the two neuroscience fields, Neurobabel also provides a quantitative link between the rodent and human neuroscience literature for exploring functional topographies across species.

## 1 Results

### 1.1 Overview

The Neurobabel framework was created in several steps. First, we built a web crawler to identify and download articles of studies that included an intracranial surgical intervention. Specifically, for this first release, we downloaded articles from ScienceDirect, if they were published after 1995 and included the keyword “bregma”, an anatomical landmark on top of the skull that is often used as reference for stereotaxic intracranial surgery. Second, we developed a text-mining tool that would extract the target coordinates of the performed surgeries. Third, we calculated the word frequency of scientific terms in each study. Specifically, we calculated the term frequency–inverse document frequency (tf-idf), which is a measure intended to reflect the importance of a word to a text document. Fourth, we combined these data with the existing Neurosynth dataset, including the word frequency of scientific terms from vocabulary shared between the included neuroimaging and behavioral neuroscience studies. This resulted in a dataset with 2668 rodent studies with 5276 surgery coordinates (Figure 1), and 6579 neuroimaging studies with 236254 activation voci, and a shared vocabulary of 2579 scientific terms.

**Figure 1:**
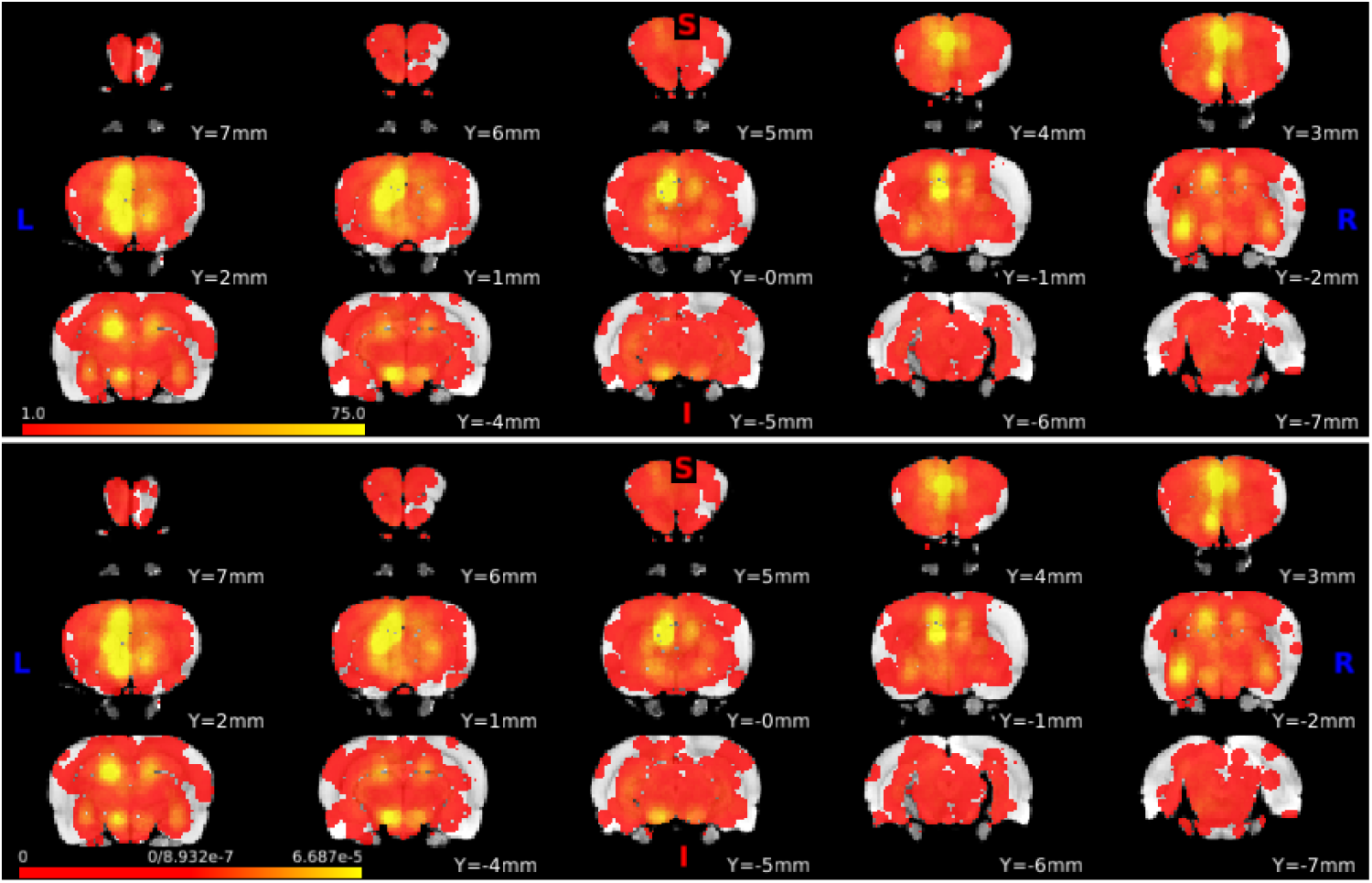
Distribution of extracted surgery coordinates. (Top) shows absolute count, in terms of number of studies, and (Bottom) shows the density, that is after dividing (Top) by the sum total of extracted surgery coordinates (n=5276). The figure shows that studies that used bregma as reference for intracranial surgery tended to focus on the rodent prelimbic and infralimbic cortex, the hippocampal region, amygdala, and hypothalamus.

We then proceeded to systematically validate the ability of our framework to provide functionality for the rodent literature analogous to that provided for the human neuroimaging literature by Neurosynth. That is, we tested whether it was possible to meta-analytically decode the function of a brain area using forward, and especially reverse inference. Next, we tested whether it was possible to use this framework to compare functional topographies across human and rodent brains.

### 1.2 Forward and Reverse Inference

One of the breakthrough functionalities of the Neurosynth framework is that it enables quantitative reverse inference, in addition to forward inference, regarding the relationship between the psychological states of participants and activation across different areas of the brain. In contrast, most hypothesis-driven neuroimaging studies depend on forward inference, which answers important questions such as, “Which brain areas become active when participants view fearful facial expressions?” Although forward inference is a powerful tool, its explanatory power falls short of that of reverse inference [12, 7], which addresses critical, conceptually distinct questions about functional specialization; for example, “Can we infer that participants were viewing fearful facial expressions, given the pattern of activation across the brain?”

Reverse inference is a powerful tool for synthesizing results across neuroimaging studies for at least two reasons. First and foremost, reverse inference takes results from studies into account that may have targeted the same brain region, but were motivated by questions from different subfields of human neuroscience. This is for example the case for the dorsal anterior cingulate region, where consensus regarding its function is only slowly emerging (for a discussion, see e.g. [9] and [13]). However, a second reason for the power of reverse inference is that neuroimaging studies often report activation in parts of the brain that were not the primary region of interest of the study. For example, a recent meta-analysis of the human striatum has reported that the anterior putamen was associated with social and language functions [11], but also that there was no single mention of the striatum in the most characteristic study for this region, in terms of striatal activation voci and decoded psychological function [14].

Reverse inference can also be used for the synthesis of results across diverse subfields of behavioral neuroscience. To demonstrate this, we compared the results of forward and reverse inferences with the psychological term “fear” from the above example. That is, forward inference asks whether it is possible to predict where surgery was performed, given the importance of the word “fear” in an article. On the other hand, reverse inference asks whether it was possible to predict whether the term fear would appear in an article, given that surgery was performed in a particular part of the brain. Given the well-established role of the amygdala in fear conditioning [15], one would predict that both forward and reverse inference would point towards this region. Furthermore, it is also well-established that the hippocampal region is involved in fear conditioning, especially in contextual fear conditioning [16]. However, because this region is involved more generally in diverse forms of memory, such as spatial and episodic memory [17, 18], reverse inference should predict a relative low probability of the word fear in an article, given that surgery was performed in this region. As shown in Fig. 2 this is exactly what we found.

**Figure 2:**
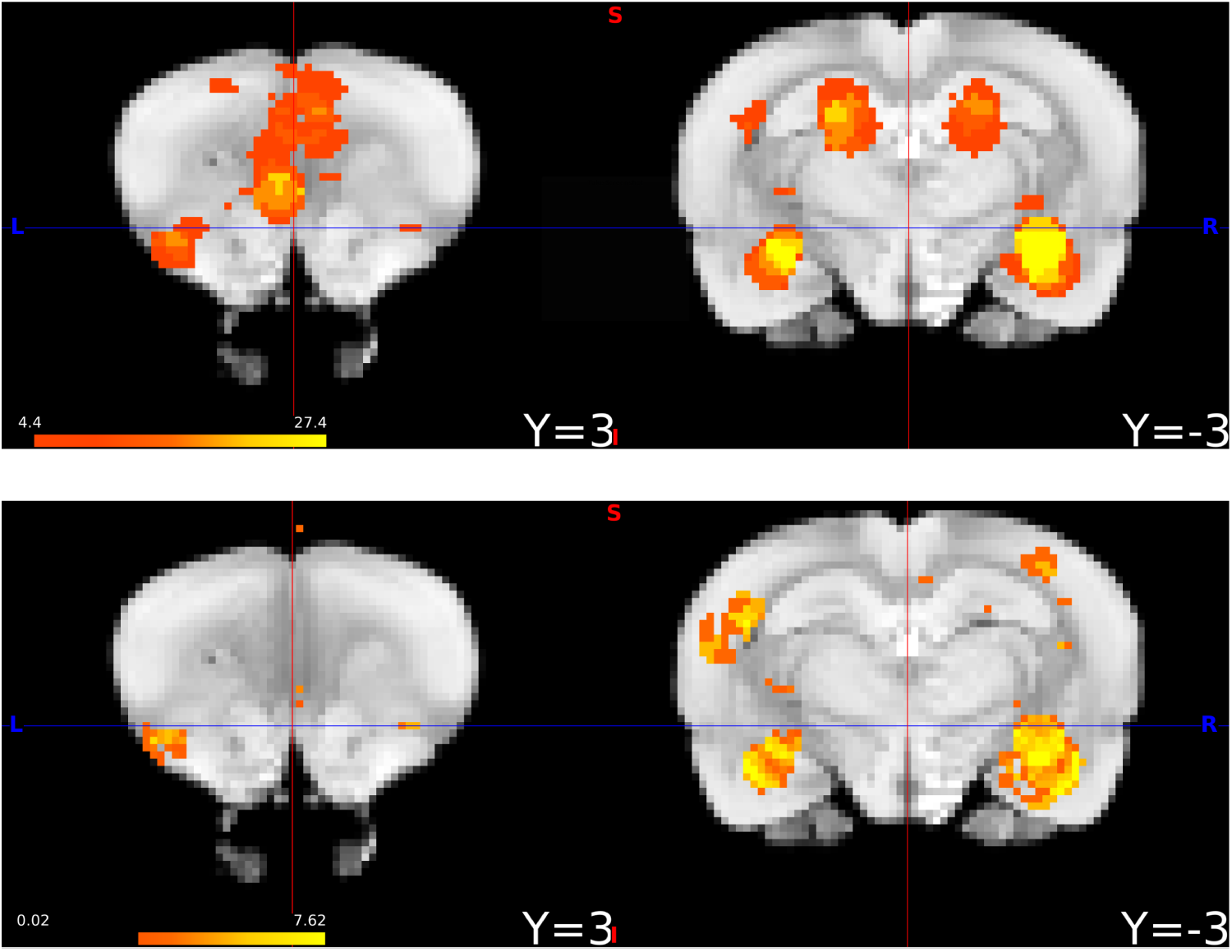
Forward and reverse inference for the term “fear”. (Top) Results (z-scores) of forward inference, that is: Given the term “fear” appeared in an article, what is the probability of intracranial surgery having been performed in each voxel of the rodent brain? (Bottom) Results of reverse inference, that is: Given that surgery was performed in a particular voxel of the rodent brain, what is probability that the term “fear” appeared in the article. One notable difference is that the hippocampal only appears during forward inference, but not during reverse inference. While this may be partially due to overall weaker results for reverse inference, it may also reflect the fact that the in comparison to the amygdala, the hippocampal formation supports spatial and contextual memory in general, while the amygdala is more specifically involved in fear-related processes. Note: For display purposes, different scales were used for forward and reverse inference results.

### 1.3 Cross-species mapping of psychological functions

The central aim for developing the present framework was to create a quantitative link between findings in behavioral neuroscience and human neuroimaging studies. This data-driven approach can be utilized to investigate similarities in functional topographies across species. This quantitative link was created in two sequential steps: First, we queried which scientific terms are associated with a specific brain region of interest in one species. We next queried which regions of the brain in the other species are associated with these terms (for more detail, see Methods subsection “Cross-species mapping of psychological functions”).

#### 1.3.1 Validation of approach

To validate our approach, we performed a test query for two well-established results in the human and animal literature. Our first query focused on well-established involvement of the amygdala in fear-related functions. Multiple studies have found that the amygdala supports the recognition of fearful facial expressions in humans [19] and during fear conditioning [16]. As a second query, we focused on the role of the hippocampus in spatial memory [17, 18].

We proceeded by first performing separate reverse inferences using the terms “fear” and “spatial memory” for rodents: For each voxel of the rodent brain, we asked: Given that surgery was performed in this voxel, what is the probability that the term appeared in the article? This resulted in a statistical map which we then projected into the human brain using the two step approach described above.

As expected, reverse inference in the rodent brain revealed a selective association of the amygdala with fear, and the hippocampal region with spatial memory (Figure 3, top left). We then applied the two-step approach for projecting these regions into the human brain. As predicted, we found that the rodent statistical map for fear was projected onto the human amygdala, while the statistical map for spatial memory was projected onto the human hippocampus (Figure 3, top right). The bottom panel of Figure 3 further shows that the top ten scientific terms linking the two species are consistent with the well-established functions of these two regions in either species. In summary, these results suggest that our approach represents a data-driven framework for investigating similarities in functional topographies across species.

**Figure 3:**
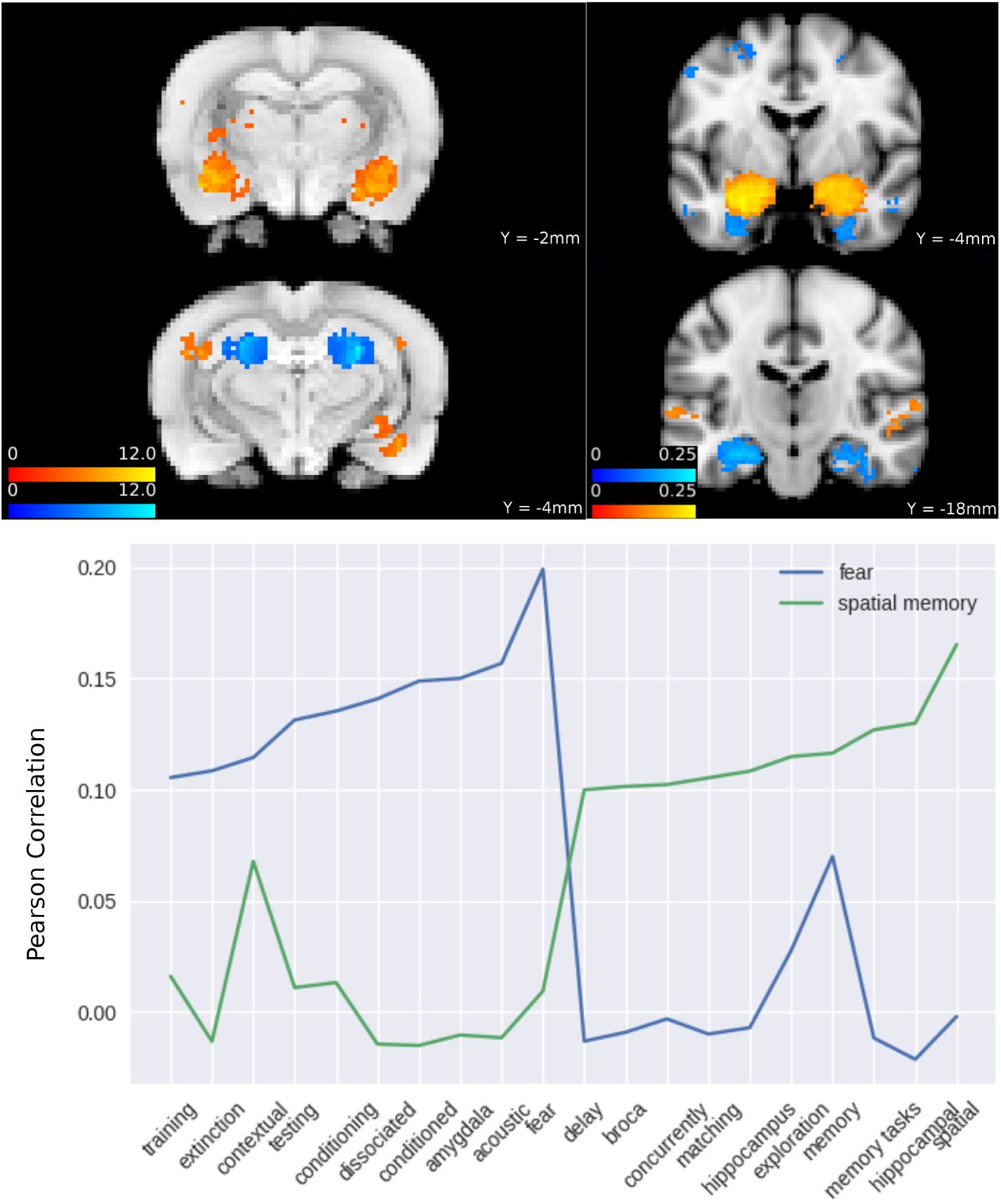
Cross-species mapping of “fear” and “spatial memory” from the rodent to the human brain. (Top-Left) Results (z-scores) of reverse inference for “fear” (orange-yellow) and “spatial memory” (blue-lightblue). (Top-Right) Human brain regions associated with the psychological terms in the center panel (Pearson correlations). (Bottom) Top-ten scientific terms associated (Pearson correlations) with brain regions identified via reverse inference for the same terms.

#### 1.3.2 Projection of rat prelimbic cortical map onto the human brain

After this validation of our approach, we proceeded to address an open question in decision neuroscience: Which human brain regions correspond to the rodent prelimbic cortex? Research in rodents has accrued a wealth of evidence that this region interacts with the dorsomedial striatum to support the acquisition and performance of goal-directed behavior [20, 21, 22]. Recent reviews of similarities in functional topographies across species have suggested that the human ventromedial prefrontal cortex may serve psychological functions similar to those functions associated with the rodent prelimbic cortex [23, 24], as the human ventromedial prefrontal cortex and the rodent prelimbic cortex seem to be similarly involved in discovering action - reward contingencies in the service of goal-directed behavior [25].

To address this question, we first created an anatomical mask for the rodent prelimbic cortex using an anatomical atlas of the rodent brain [26]. We then used the above two-step approach to project this region into the human brain. We found that this resulted in a distributed set of clusters in the human brain, including the lateral prefrontal cortex, anterior amygdala, and dorsomedial prefrontal cortex (Figure 4, top). Interestingly, despite this distributed set of brain regions, we found that the top ten scientific terms associated with the rodent prelimbic cortex were predominantly anatomical terms referring to the medial prefrontal cortex 4, bottom).

**Figure 4:**
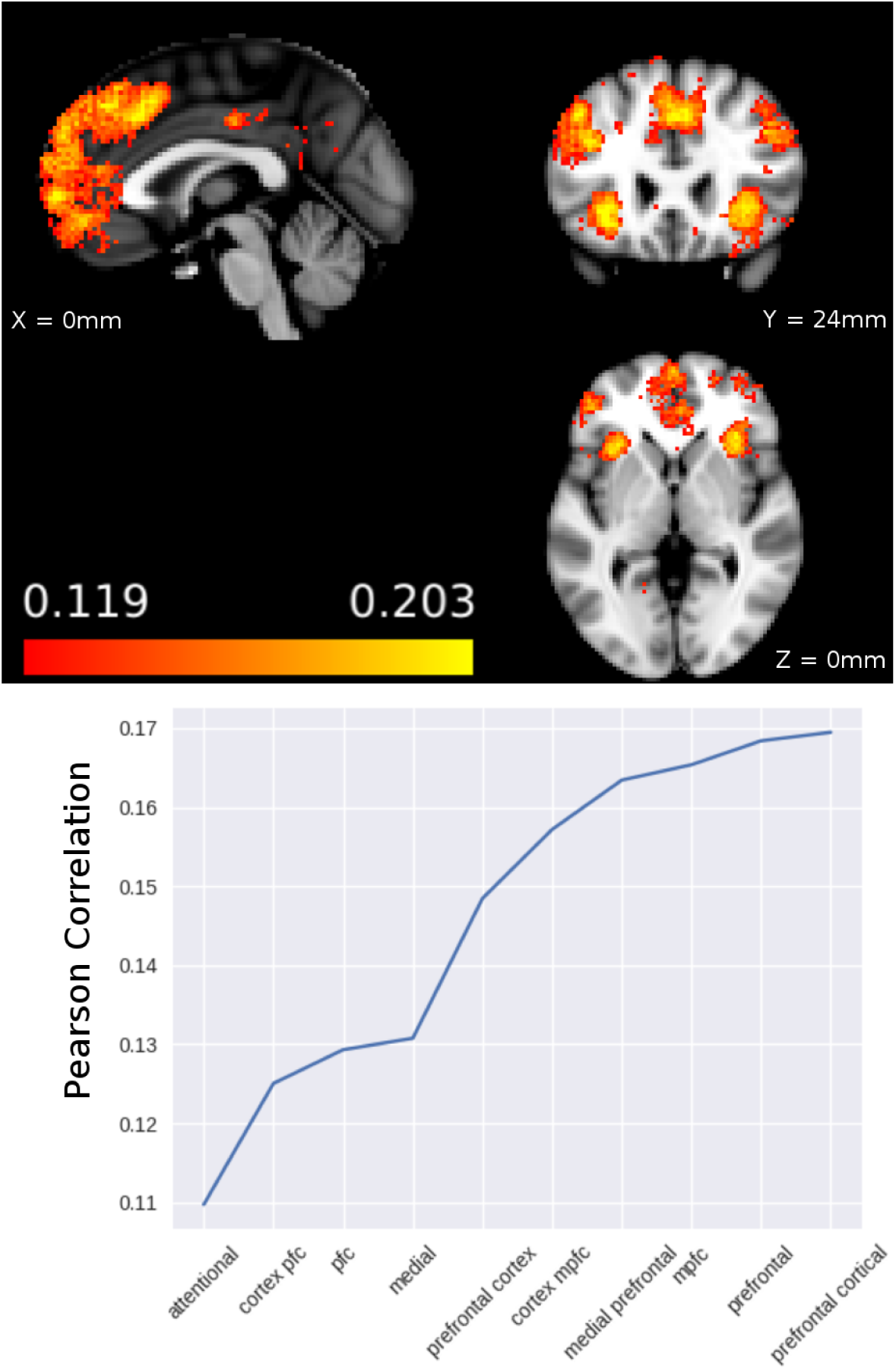
Cross-species mapping of the rodent prelimbic cortex to the human brain. (Top) The cross-species mapping identified a distributed set of clusters, including the medial dorsal and ventral frontal cortices, dorsolateral prefrontal cortex, as well as anterior insular cortex. Overlay shows Pearson correlations. (Bottom) Top-ten scientific terms identified (Pearson correlation) in the decoding of the anatomical region of the prelimbic cortex in rodents.

#### 1.3.3 Projection of human dorsolateral prefrontal cortex onto the rat brain

Next, we utilized our framework to query it for the rodent pendant of the human dorsolateral prefrontal cortex. Whether rodents have a pendant of the human dorsolateral prefrontal cortex is still a matter of debate. It has been argued that based on anatomical and functional characteristics, the rodent prelimbic cortex, dorsal anterior cingulate cortex, and frontal cortical area 2 may constitute a rodent pendant of dorsolateral prefrontal cortex (for a review, see [27]). To address this questions, we followed an approach analogous to the above query regarding the human pendant of the rodent prelimbic cortex. We created an anatomical mask of the middle frontal gyrus, according to the Harvard-Oxford human anatomical atlas [28, 29]. Next, we decoded the psychological function of this area in humans to investigate which regions in the rodent brain are associated with these psychological functions. We found that this procedure lead to an almost exclusive selection of the rodent prelimbic cortex and the anterior cingulate cortex (see Figure 5, top). The bottom panel of figure 5 shows the top ten scientific terms associated with the anatomical mask of the middle frontal gyrus in humans.

**Figure 5:**
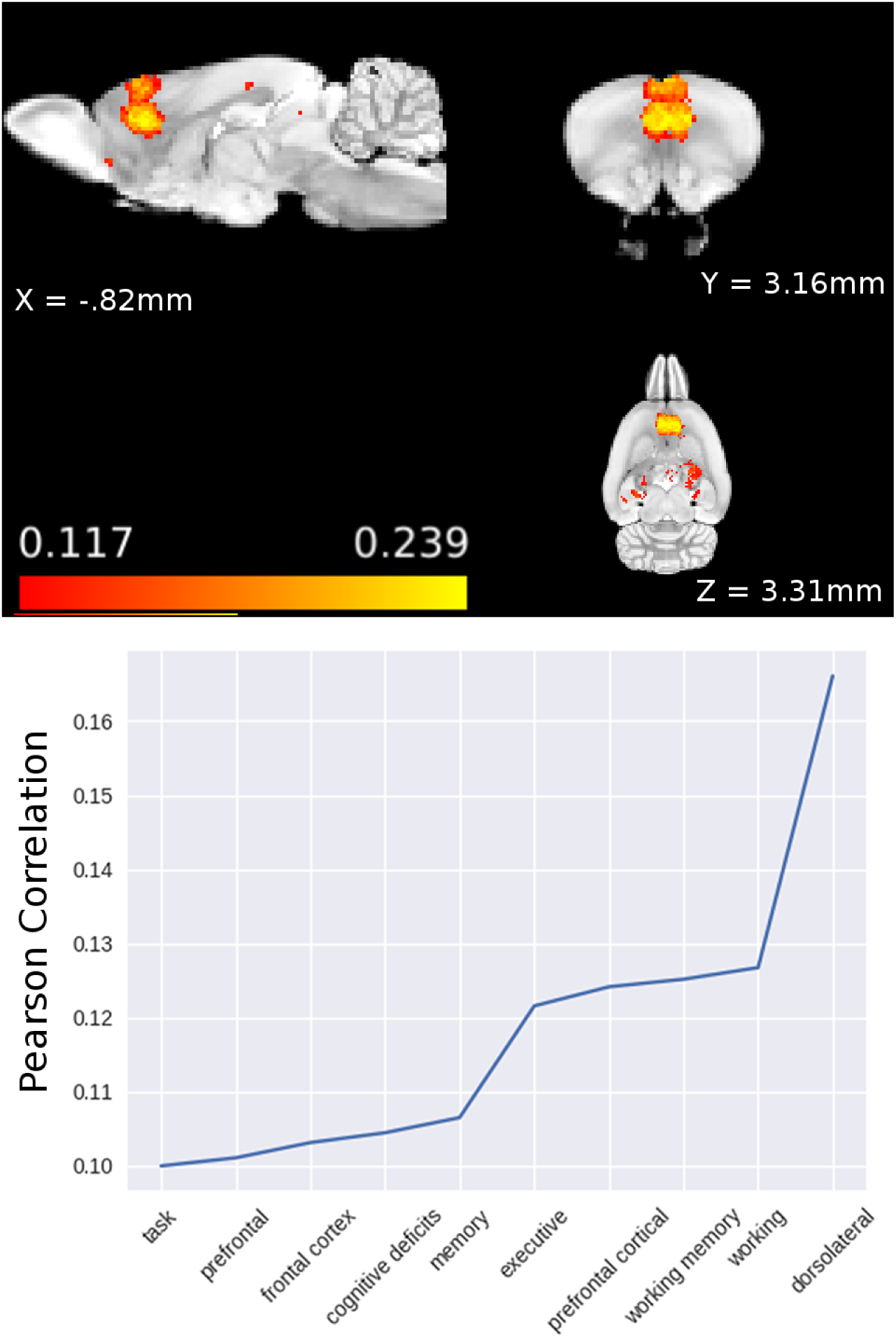
Cross-species mapping of the human medial frontal gyrus into the rodent brain. (Top) The cross-species mapping identified the rodent prelimbic and anterior cingulate cortex to correspond to the human medial frontal gyrus. Overlay shows Pearson correlations. (Bottom) Top-ten scientific terms associated (Pearson correlations) with an anatomical map of the human medial frontal gyrus.

## 2 Discussion

The aim of this work was to create an easily expandable extension of the well-established Neurosynth framework for human neuroimaging data, to provide (1) analogous analytic tools for the rodent literature, and (2) a data-driven tool for characterizing similarities of functional topographies in human and rodent brains.

When we applied our semantic functional cross-species mapping procedure to identify the human functional pendant of the rodent prelimbic cortex, the results was distributed set of frontal cortical regions in humans. This result is interesting in the context of the emerging view in the field that the human ventromedial prefrontal cortex shares functional similarities with the rodent prelimbic cortex [23]. While a previous review [27] suggested that the medial surface of the rodent frontal cortex is more differentiated in humans, and distributed across the lateral prefrontal cortex, our results go beyond this by including the anterior insular cortex. Future analyses may be able to disentangle whether this inclusion of the anterior insular cortex is caused simply by differences in the functional architectures across species, or whether it is a fundamental short-coming of the present approach, which relies on how scientist describe the methods, results, and aims of their studies. At the same time, consistent with our findings, a previous meta-analysis with Neurosynth reported that the anterior insular cortex interacted with frontal cortical regions in support of cognitive functions [8].

When we conversely queried for the functional pendant of the human lateral prefrontal cortex in rodents, we found an almost exclusive selection of the prelimbic cortex and dorsal anterior cingulate cortex. This result is consistent with the above finding, where the rodent prelimbic cortex was functionally mapped onto a distributed set of frontal cortical regions, including the lateral prefrontal cortex.

Several critical limitations of our dataset and approach should be kept in mind. Some of these are the same as those of the original Neurosynth framework. Most critically, the results depend largely on the consensus in the field regarding the function of a brain region. For example, consider the case that only a small number of studies have uncovered the true psychological function of a brain region, while a large number of studies assign a different, potentially less specific or less appropriate, function to the same brain region. In this case, the results from applying the present approach would most likely be dominated by the large number of studies that reached an inaccurate consensus regarding the psychological function of this region, rather than by the small number of studies which have uncovered the true psychological function of a brain region.

A further limitation of the present dataset is that it only includes surgery coordinates with the anatomical landmark bregma as reference. This limitation can be easily overcome in future data releases by incorporating studies that used surgery coordinates with other anatomical references (e.g. interaural). However, this limitation is substantial, and should be kept in mind when interpreting the results of forward and reverse inference, as well as of the semantic functional cross-species mapping procedure.

Related to the previous limitation, the present dataset is limited by where in the brain researchers have performed intracranial surgery. That is, the Neurobabel framework will not be able to support the discovery of functions in brain regions that have not been targeted with intracranial surgery. This limitation is largely absent for the original Neurosynth dataset, because neuroimaging studies often collect functional data from the whole brain, even if the aim of a research project is focused on a particular part of the brain.

This difference between the present dataset for behavioral neuroscience studies and the original dataset for neuroimaging is also important when interpreting diverging results of forward and reverse inference in rodents. In the original Neurosynth dataset, this divergence is driven by two factors. First, by the fact that reverse inference is relatively more apt at integrating findings across studies from different neuroimaging subfields, which may investigate the role of a particular brain region across a diverse set of psychological functions. Second, human neuroimaging studies oftentimes include tables reporting activation coordinates in parts of the brain that were not the focus of a study.

Both these factors contribute to the strength of reverse inference as provided by the Neurosynth framework. In contrast, in the present dataset only the first of the two factors contributes to different results from forward relative to reverse inference.

We consider the present results a demonstration of some of the capabilities provided by the current dataset. Several extension and improvements are plausible and easily attainable.

Similar to how the original Neurosynth dataset has grown in size since its initial conception, the automated nature of aggregating studies, extracting surgery coordinates, and text-mining of companion articles will also allow the present dataset to evolve over time. Furthermore, as mentioned above, future release should also consider surgery coordinates that are not reported with bregma as reference.

Beyond the demonstrations of the data-driven approach presented here, the dataset will enable scientists to ask various other research questions. For example, this dataset could be used by scientists from one field of neuroscience to determine which scientists in the other field may be interested in collaborations. Another application of this dataset could be to not only consider where surgery was performed, but also which manipulation (e.g. micro-infusion versus lesion), or which pharmacological manipulation was performed.

Overall, the results presented here demonstrate the feasibility of a natural extension of the Neurosynth framework to also include rodent studies. We believe that this approach could lead to increased collaboration between the fields of human and behavioral neuroscience. In particular, it may facilitate efforts of determining to which degree findings from animal models may translate into human neuroscience. This could lead to the development of novel targeted psychiatric interventions. All data and code have been made open-source, to facilitate future collaboration.

## 3 Methods

### 3.1 Automated coordinate extraction

Surgery coordinates were extracted algorithmically, by first identifying the methods section of an article, followed by an identification of sentences which included the keyword “bregma”. Bregma is an anatomical landmark, which is frequently used as reference during intracranial surgeries. Bregma is the anatomical point on the skull at which the coronal suture is intersected perpendicularly by the sagittal suture. Coordinates were then extracted using regular expressions, working under the assumption that coordinates would be expressed in numbers between the keywords “anterior” or “posterior”, “medial” or “lateral”, and “dorsal” or “ventral”. An ordered sequence of more relaxed regular expression was tested consecutively, until a match was found. Details are provided in the github repository for this project: https://github.com/wmpauli/ACE/.

The present dataset does not take into account the age or weight of rodents included in each of the studies. While this should ideally be done, to achieve a higher accuracy for surgery coordinates, we argue that due to the large number of included studies, the effect of age and weight of rodents should not introduce a systematic bias.

### 3.2 Correction for different habits of performing surgery

In the behavioral neuroscience literature, surgery coordinates are sometimes expressed in distance from dura or skull, and sometimes with bregma as dorsal/ventral reference. After coordinate extraction, we therefore identified studies in which the depth of the surgical intervention was expressed in distance from dura or skull. For these studies, we applied a non-linear correction of the reported coordinates to take into account the medial-lateral and anterior-posterior curvature of the surface of the rodent skull. The amount of adjustment was determined using an anatomical atlas [26].

### 3.3 Data Preparation

After extraction of surgery coordinates, coordinates where converted to the Waxholm Space Atlas of the Sprague Dawley Rat Brain [30], to enable the application of standard neuroimaging libraries for data analysis. A 1mm boxcar smoothing kernel was applied to surgery coordinates.

### 3.4 Vocabulary of psychological terms

To create a vocabulary shared between behavioral neuroscience and neuroimaging studies, we included psychological terms that were included in the 2015 data release of neurosynth (version 0.6) and also occurred in the behavioral neuroscience studies included in the present dataset (term frequency–inverse document frequency (tf-idf) threshold of 0.001). A total of 2579 psychological terms was included.

### 3.5 Anatomical ROIs

Anatomical regions of interest (ROIs) were determined according to an anatomical atlas [26].

### 3.6 Forward and Reverse inference

The present extension of the neurosynth dataset was developed with the explicit intent that the all of the existing analysis tools of the neurosynth framework could be leveraged to perform analogous forward and reverse inference analyses, as has been done successfully for neuroimaging data [7]. We refer to the original publication and neurosynth.org for details of the approach. Briefly, for forward inference we calculated the probability that surgery was performed at a given coordinate, given that a psychological term appeared in the companion article (i.e. *P* (*xyz*|*term*)). For reverse inference, we calculated *P* (*term*|*xyz*). We present the results of these analyses using the standard approach of neurosynth, i.e. z-scores reflecting the probabilities. For display purposes we applied a uncorrected threshold of p=0.001 to statistical maps.

### 3.7 Cross-species mapping of psychological function

To perform cross-species mappings, we decided to follow an approach that entirely relies on Pearson correlations. We made this choice for two reasons: (1) consistency of this analysis approach with existing methods implemented in the neurosynth framework (“decode” method), and (2) because of the intuitive understanding of Pearson correlation by a broad readership. More, advanced machine learning techniques could be applied to the present dataset as well. For example, rather than relying simply on the word frequency (tf-idf) of scientific terms, the present dataset would allow the application of more advanced natural language processing algorithms.

The cross-species mapping was performed in two steps. First, we decoded the psychological function of each voxel, by calculating the Pearson correlation between activity/surgery in each voxel and the occurrence (tf-idf) of each scientific term across studies. This was done separately for each species, yielding two (one per species) *v*_*s*_ x *t* voxel-by-term matrices *X*_*s*_, where *v*_*s*_ is the number of voxels in a species *s*, and *t* is the number of scientific terms in the vocabulary shared between human and animal literature (this step was performed using the “decode” function of the neurosynth framework). To perform the functional cross-species mapping of a brain region of interest in one species to the other species, we first normalized each matrix *X*_*s*_, by subtracting the row-means, and then dividing by the square-root of row-sum of squares. This allowed us to calculate Pearson correlations with dot products of matrices. We decoded the function of a region of interest by calculating the dot product of the associated 1 x *v*_*s*_ brain mask *M*_*s*_ and *X*_*s*_, yielding a 1 x *t* term vector *T*. Next, we determined which brain voxels in the other species were associated with this term vector by calculating the dot product of the (normalized) term vector *T* and the transpose of the voxel-by-term matrix *X*_*s*_ of the other species. This yielded a 1 x *v*_*s*_ vector indicating how strongly the function of each voxel in the second species correlated with the function of the region of interest in the first species. In addition, this analysis produced the term vector *T*, representing a functional/quantitative link between the two statistical maps.

### 3.8 Code Availability

The source code for creating the dataset and performing the analyses reported here are available as a github repository: https://github.com/wmpauli/neurosynth.

### 3.9 Data Availability

The present release of the dataset is available in the following github repository: https://github.com/wmpauli/neurosynth-data. The cross-species mapping of psychological functions depends on what is referred to as feature images in the Neurosynth framework. Because the process of creating these feature images takes a many hours, we uploaded the output of this process to the open science framework (OSF): DOI 10.17605/OSF.IO/6D3B2.

## 4 Acknowledgments

Thank you to Dr. Jane E. Barker and Dr. Julian M. Tyszka for constructive discussions of the approach, and for proofreading the manuscript. Further thanks go to Dr. Tal Yarkoni for opening up the source code for neurosynth to the public.

## Notes

#### Summary of Updates

In response to comments from other researchers, I made some small changes to wording.

## References

[1] Everitt, B. J. & Robbins, T. W. Drug Addiction: Updating Actions to Habits to Compulsions Ten Years On. Annual Review of Psychology 67, 23–50 (2016). URL http://dx.doi.org/10.1146/annurev-psych-122414-033457.

[2] Mulholland, P. J., Chandler, L. J. & Kalivas, P. W. Signals from the Fourth Dimension Regulate Drug Relapse. Trends in Neurosciences 39, 472–485 (2016). URL http://www.sciencedirect.com/science/article/pii/S0166223616300170.

[3] EMCDDA. European Drug Report 2016: Trends and Developments (2016). URL http://www.emcdda.europa.eu/publications/edr/trends-developments/2016_en.

[4] NIDA. Trends & Statistics (2017). URL https://www.drugabuse.gov/related-topics/trends-statistics.

[5] Hyman, S. E. Revolution Stalled. Science Translational Medicine 4, 155cm11–155cm11 (2012). URL http://stm.sciencemag.org/content/4/155/155cm11.

[6] Cressey, D. Psychopharmacology in crisis. Nature 201110 (2011).

[7] Yarkoni, T., Poldrack, R. A., Nichols, T. E., Van Essen, D. C. & Wager, T. D. Large-scale automated synthesis of human functional neuroimaging data. Nature Methods 8, 665–670 (2011). URL http://www.nature.com/nmeth/journal/v8/n8/full/nmeth.1635.html.

[8] Chang, L. J., Yarkoni, T., Khaw, M. W. & Sanfey, A. G. Decoding the Role of the Insula in Human Cognition: Functional Parcellation and Large-Scale Reverse Inference. Cerebral Cortex 23, 739–749 (2013). URL http://cercor.oxfordjournals.org/content/23/3/739.

[9] Lieberman, M. D. & Eisenberger, N. I. The dorsal anterior cingulate cortex is selective for pain: Results from large-scale reverse inference. Proceedings of the National Academy of Sciences 112, 15250–15255 (2015). URL http://www.pnas.org/content/112/49/15250.

[10] Vega, A. d. l., Chang, L. J., Banich, M. T., Wager, T. D. & Yarkoni, T. Large-Scale Meta-Analysis of Human Medial Frontal Cortex Reveals Tripartite Functional Organization. Journal of Neuroscience 36, 6553–6562 (2016). URL http://www.jneurosci.org/content/36/24/6553.

[11] Pauli, W. M., O’Reilly, R. C., Yarkoni, T. & Wager, T. D. Regional specialization within the human striatum for diverse psychological functions. Proceedings of the National Academy of Sciences 113, 1907–1912 (2016). URL http://www.pnas.org/content/113/7/1907.

[12] Poldrack, R. A. Can cognitive processes be inferred from neuroimaging data? Trends in Cognitive Sciences 10, 59–63 (2006). URL http://www.sciencedirect.com/science/article/pii/S1364661305003360.

[13] Wager, T. D. et al. Pain in the ACC? Proceedings of the National Academy of Sciences 113, E2474–E2475 (2016). URL http://www.pnas.org/content/113/18/E2474.

[14] Zarate, J. M. & Zatorre, R. J. Experience-dependent neural substrates involved in vocal pitch regulation during singing. NeuroImage 40, 1871–1887 (2008). URL http://www.sciencedirect.com/science/article/pii/S1053811908000591.

[15] LeDoux, J. E., Iwata, J., Cicchetti, P. & Reis, D. J. Different projections of the central amygdaloid nucleus mediate autonomic and behavioral correlates of conditioned fear. The Journal of Neuroscience 8, 2517–2529 (1988). URL http://www.jneurosci.org/content/8/7/2517.

[16] Phillips, R. G. & LeDoux, J. E. Differential contribution of amygdala and hippocampus to cued and contextual fear conditioning. Behavioral Neuroscience 106, 274–285 (1992).

[17] O’Keefe, J. & Dostrovsky, J. The hippocampus as a spatial map. Preliminary evidence from unit activity in the freely-moving rat. Brain Research 34, 171–175 (1971). URL http://www.sciencedirect.com/science/article/pii/0006899371903581.

[18] Eichenbaum, H., Dudchenko, P., Wood, E., Shapiro, M. & Tanila, H. The Hippocampus, Memory, and Place Cells: Is It Spatial Memory or a Memory Space? Neuron 23, 209–226 (1999). URL http://www.sciencedirect.com/science/article/pii/S0896627300807734.

[19] Adolphs, R., Tranel, D., Damasio, H. & Damasio, A. R. Fear and the human amygdala. Journal of Neuroscience 15, 5879–5891 (1995). URL http://www.jneurosci.org/content/15/9/5879.

[20] Baker, P. M. & Ragozzino, M. E. Contralateral disconnection of the rat prelimbic cortex and dorsomedial striatum impairs cue-guided behavioral switching. Learning & Memory 21, 368–379 (2014). URL http://learnmem.cshlp.org/content/21/8/368.

[21] Balleine, B. W. & Dickinson, A. Goal-directed instrumental action: contingency and incentive learning and their cortical substrates. Neuropharmacology 37, 407–419 (1998). URL http://www.sciencedirect.com/science/article/pii/S0028390898000331.

[22] Ragozzino, M. E., Ragozzino, K. E., Y, J. & Kesner, R. P. Role of the dorsomedial striatum in behavioral flexibility for response and visual cue discrimination learning. Behavioral Neuroscience 116, 105–115 (2002).

[23] Balleine, B. W. & O’Doherty, J. P. Human and rodent homologies in action control: corticostriatal determinants of goal-directed and habitual action. Neuropsychopharmacology 35, 48–69 (2009). URL http://www.nature.com/npp/journal/v35/n1/full/npp2009131a.html.

[24] O’Doherty, J. P., Cockburn, J. & Pauli, W. M. Learning, reward, and decision making. Annual Review of Psychology 68, 73–100 (2017).

[25] Liljeholm, M., Tricomi, E., O’Doherty, J. P. & Balleine, B. W. Neural Correlates of Instrumental Contingency Learning: Differential Effects of Action–Reward Conjunction and Disjunction. Journal of Neuroscience 31, 2474–2480 (2011). URL http://www.jneurosci.org/content/31/7/2474.

[26] Paxinos, G. & Watson, C. A stereotaxic atlas of the rat brain. New York: Academic (1998).

[27] Uylings, H. B. M., Groenewegen, H. J. & Kolb, B. Do rats have a prefrontal cortex? Behavioural Brain Research 146, 3–17 (2003). URL http://www.sciencedirect.com/science/article/pii/S0166432803003346.

[28] Desikan, R. S. et al. An automated labeling system for subdividing the human cerebral cortex on MRI scans into gyral based regions of interest. NeuroImage 31, 968–980 (2006).

[29] Frazier, J. A. et al. Structural brain magnetic resonance imaging of limbic and thalamic volumes in pediatric bipolar disorder. The American Journal of Psychiatry 162, 1256–1265 (2005).

[30] Papp, E. A., Leergaard, T. B., Calabrese, E., Johnson, G. A. & Bjaalie, J. G. Waxholm Space atlas of the Sprague Dawley rat brain. NeuroImage 97, 374–386 (2014). URL http://www.sciencedirect.com/science/article/pii/S1053811914002419.

